# Surface texture guides egg-laying decisions in *Aedes aegypti* mosquitoes

**DOI:** 10.64898/2026.03.08.710377

**Authors:** Alexandra Anoshina, Nicholas K. Tochor, Lauren Semkow, Annie Zeng, Benjamin J. Matthews

## Abstract

Mosquitoes undergo development as aquatic larvae and pupae before emerging as terrestrial adults. Accordingly, blood-fed and mated female mosquitoes must select an appropriate egg-laying site to maximize the fitness of their offspring. Female yellow fever mosquitoes (*Aedes aegypti*) lay their eggs above the waterline of small containers or natural bodies of water, where they can remain dormant for many months until they are submerged and hatch. Here, we show that female mosquitoes use surface texture as a powerful cue to guide egg-laying decisions, selecting rougher textures over smooth when choosing among containers and when selecting specific sites within a given substrate. In addition, we identify an interaction between substrate texture and water salinity with respect to egg-laying decisions, demonstrating that female mosquitoes integrate competing cues to determine the ultimate suitability of an egg-laying site. Finally, we explore the dynamics of local egg-laying search behaviour, demonstrating that texture modulates traversal speed while mosquitoes search for appropriate egg-laying sites.

## Introduction

Selecting a suitable egg-laying site is critical to fitness, particularly for mosquitoes and other insects that undergo early development in aquatic environments. The dangers associated with inappropriate site selection include a lack of food for developing larvae, the presence of predators or other natural enemies, exposure to lethal temperatures or salinities, and the disappearance of water altogether. Female yellow fever mosquitoes (*Aedes aegypti*) lay eggs in clutches of 100-150 near a variety of natural (e.g. tree holes, rock depressions) or artificial (e.g. tires, water storage containers) bodies of water (*1–3*). They lay eggs above the water line, where they form a hard scleratized and melanized outer shell to prevent dessication. Following this process the eggs can remain viable for months to years until submerged in water (*3, 4*). When inundated, the eggs hatch into aquatic larvae that develop via filter feeding over the course of days to weeks. This strategy is thought to have arisen in their ancestral sub-Saharan range to prevent premature hatching in quickly dwindling water sources as the dry season approaches (*5, 6*). As eggs can remain viable for up to 6 months, *Ae. aegypti* are a particularly pernicious and well-adapted disease vector in areas of changing climate and fluctuating precipitation (*7, 8*).

To select a suitable egg-laying site, mosquitoes rely on a variety of long-, mid-, and short-range sensory cues (*1*). These cues act on an array of sensory modalities that can operate at a distance, such as hygrosensation (*9, 10*), olfaction, and vision. Once they have identified a potential egg-laying site from a distance, female mosquitoes use contact-mediated gustatory and mechanosensory cues to generate a final decision about whether to lay an egg (*11–13*).

Surface texture is a known egg-laying cue for insects broadly (*14, 15*), including for Aedine mosquitoes (*16–21*). For example, Fay and Perry found that *Ae. aegypti* females show a significantly stronger preference for brown blotting paper and brown bag papers over white filter paper or aluminum foil (*20*). In another field-based study, Chadee et al. demonstrated that female *Ae. aegypti* prefer to lay eggs on rough substrates which they suggested may facilitate resting and egg-laying (*19*). Finally, O’Gower found that *Ae. aegypti* used a variety of chemical, visual, and tactile cues to guide egg-laying, with tactile cues often dominating over other modalities when free water is available (*13, 22*). Despite these studies, carefully controlled studies to assess how texture guides egg-laying and interacts with other egg-laying cues remain limited.

To better understand the role of mechanical cues in *Ae. aegypti* egg-laying behaviour we first sought to characterize the impact of surface texture (roughness) in isolation from other egg-laying cues in controlled laboratory assays. We hypothesized that altered substrate texture would impact egg-laying behaviour and preference, predicting that rougher textures would be preferred over smooth, and that female mosquitoes would employ search strategies that maximized the likelihood that they would lay eggs on rough surfaces. To test this, we performed a series of two-choice preference assays at different spatial scales to determine how different textures influenced egg-laying preference with respect to site selection both between and within individual containers. We explored differences in how eggs were deposited relative to one another depending on whether eggs were deposited on rough or smooth surfaces and investigated the interaction between rough texture (a positive cue) with elevated salinity (a negative cue). Finally, we performed video analysis to determine how surface roughness influences mosquito locomotion during egg-laying bouts.

## Results

### Rough substrate texture is a positive egg-laying cue

To test texture preference during egg-laying, we created a two-choice egg-laying assay where ten blood-fed and mated female mosquitoes were placed in a small chamber with access to two petri dishes, each with a small pool of water and an agarose egg-laying substrate of varied surface textures (Figure 1A). *Ae. aegypti* showed an almost universal preference for all textured substrates over a smooth alternative. The strength of this preference was correlated with the degree of roughness of the substrate, i.e. it increased with the average particle size of the substrate surface (Figure 1B). The relationship between egg-laying preference and surface texture can be described with a log-logistic function (Figure 1C and see Methods). Analysis of egg numbers revealed a weakly negative relationship between the identity of the textured choice and total egg number across both plates of the assay (Figure 1D).

**Figure 1.**
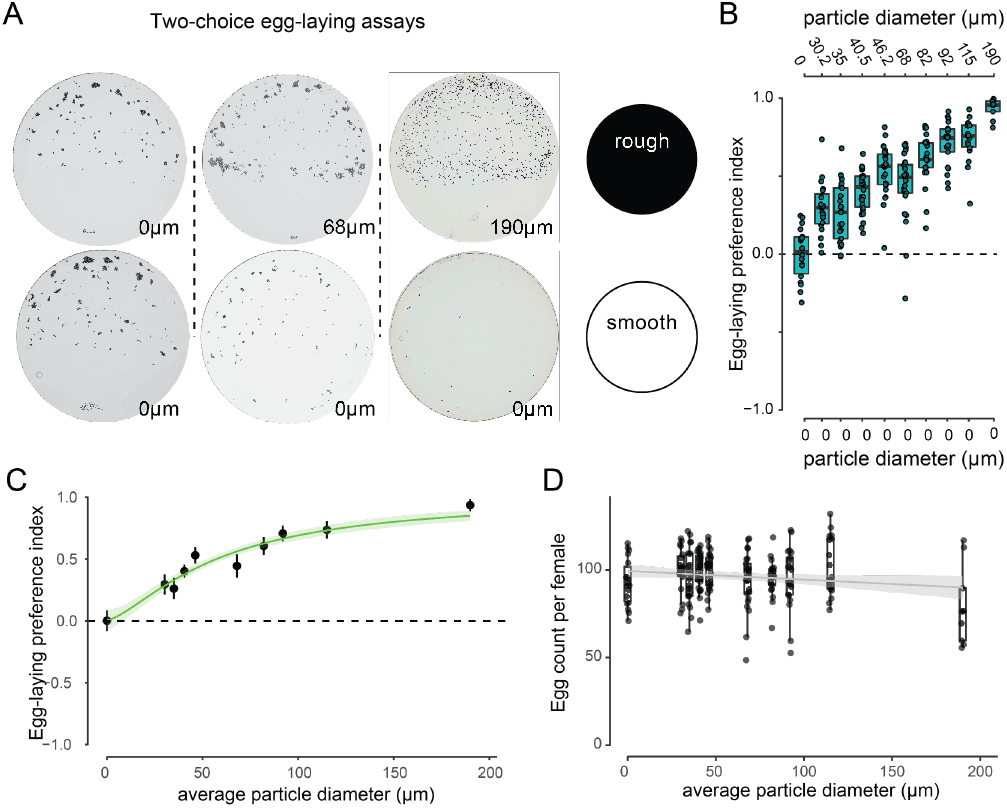
Mosquitoes prefer rough egg-laying substrates. **(A)** Representative images of *Ae. aegypti* eggs deposited on 60mm agarose plates imprinted with sandpaper of different average particle diameters (two-choice assays with textures indicated, left to right: 0µm vs.0µm, 68µm vs. 0µm, and 190µm vs 0µm). **(B)** Egg-laying preference index (PI) measured in two-choice assays in which females chose between a smooth agarose substrate and an agarose substrate textured with sand-paper of increasing average particle diameter. A positive PI indicates preference for the textured substrate, and each assay consisted of 10 blood-fed mosquitoes. A one-way ANOVA followed by Tukey HSD pairwise tests indicates that all textured surfaces show increased preference relative to control. **(C)** Log-logistic function (fixed upper bound of 1 and lower bound of 0) fit to the egg-laying preference data shown in B (slope (b) = −1.4394; ED50 (e) = 57.20µm). **(D)** Total number of eggs laid per female in two-choice assays (smooth vs. plates with the indicated textures on the x-axis) fit with a linear regression (slope = −.04960, adjusted R2 = 0.01453, p=0.047). n=9-26 assays per condition.

### Texture guides egg-laying decisions at small scale

To determine the spatial scale at which texture preference acts, we created a variant of our two-choice assay in which textured alternatives are directly adjacent to one another in a ‘split-dish’ format (Figure 2A). When we modulated the texture to include textures of different roughness on each side, we observed a preference for the half of the dish that contained the rougher texture (Figure 2B) which tracked with the relative difference in average particle diameter (Figure 2C). Interestingly, the model fit to the relative texture data is qualitatively similar to the model fit to the two-choice assays (Figure 1C), suggesting that mosquitoes can maintain similar relative values while choosing substrate both within and between containers.

**Figure 2.**
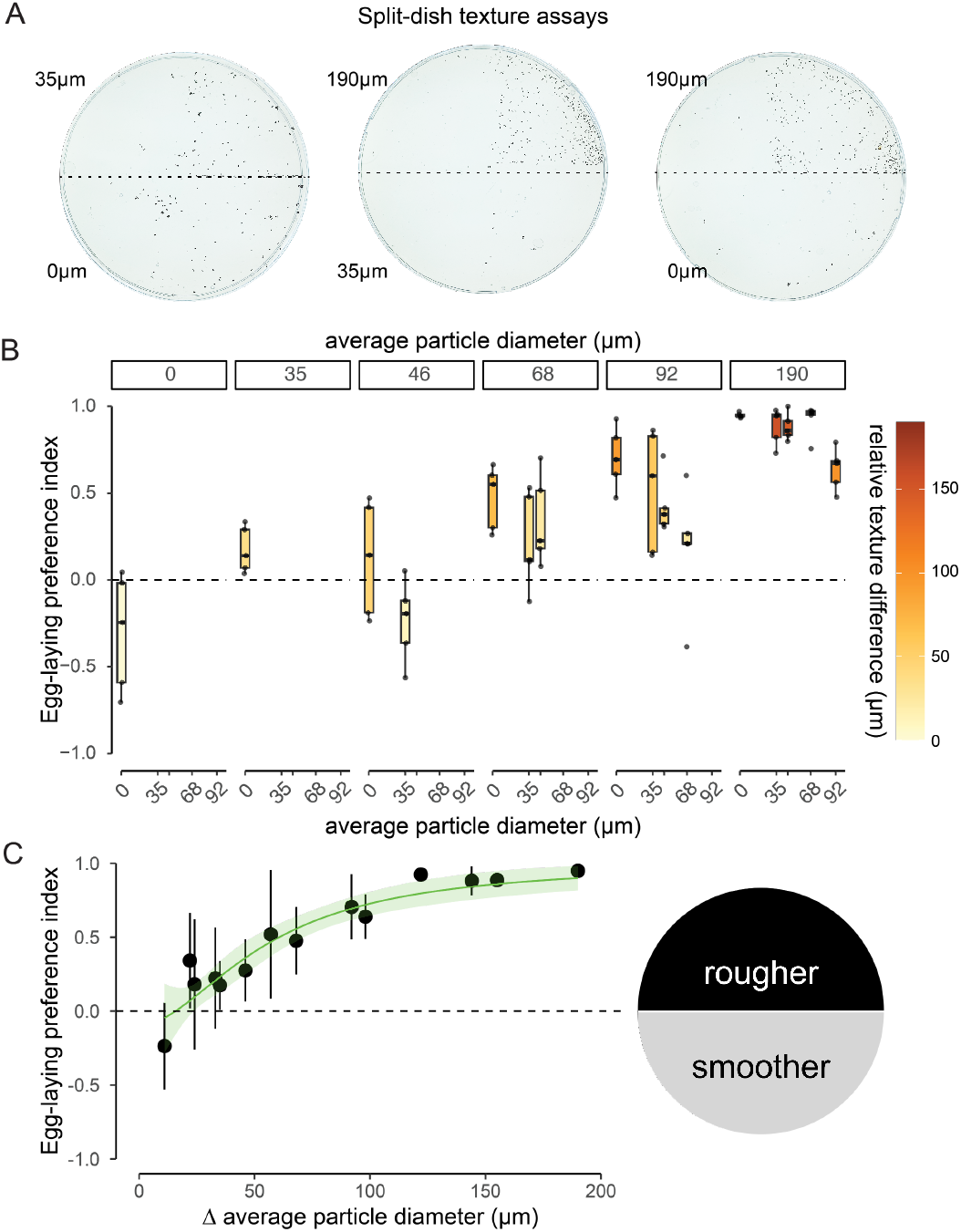
Rough texture preference is maintained at small scales. **(A)** Representative images of split Petri dishes used to assay egg-laying preference between substrates differing in particle size. Each dish contained two halves, each patterned with a texture with a defined average particle diameter as indicated. **(B)** Egg-laying preference indices of mosquitoes offered split-petri dishes with the indicated textures. For each boxplot, the relative texture difference of the two halves is represented by the colour scale at right. **(C)** Log-logistic function (fixed upper bound of 1 and lower bound of 0) fit to the egg-laying preference data shown in B (slope (b) = −2.169; ED50 (e) = 62.77µm. Note the similarities to the function from Figure 1C. n=5-10 per condition.

### Eggs are ‘clumped’ on smooth surfaces

In instances where mosquitoes lay eggs on smooth surfaces, they appear to ‘clump’ their eggs together as opposed to evenly spread them across the surface (Figure 3A). To quantify the degree of egg clustering, we calculated the distance of each eggs’ closest neighbors. We found that at rougher textures eggs were more likely to be distributed singly at rougher textures (Figure 3B). The tendency to lay eggs in ‘clumps’ at smoother textures was demonstrated by analyzing the mean distance between an egg and its six nearest neighbors (Figure 3C). We note that the data are heavily skewed, with peak average distances at the short end and few points at the long end representing single eggs scattered around the larger egg clusters.

**Figure 3.**
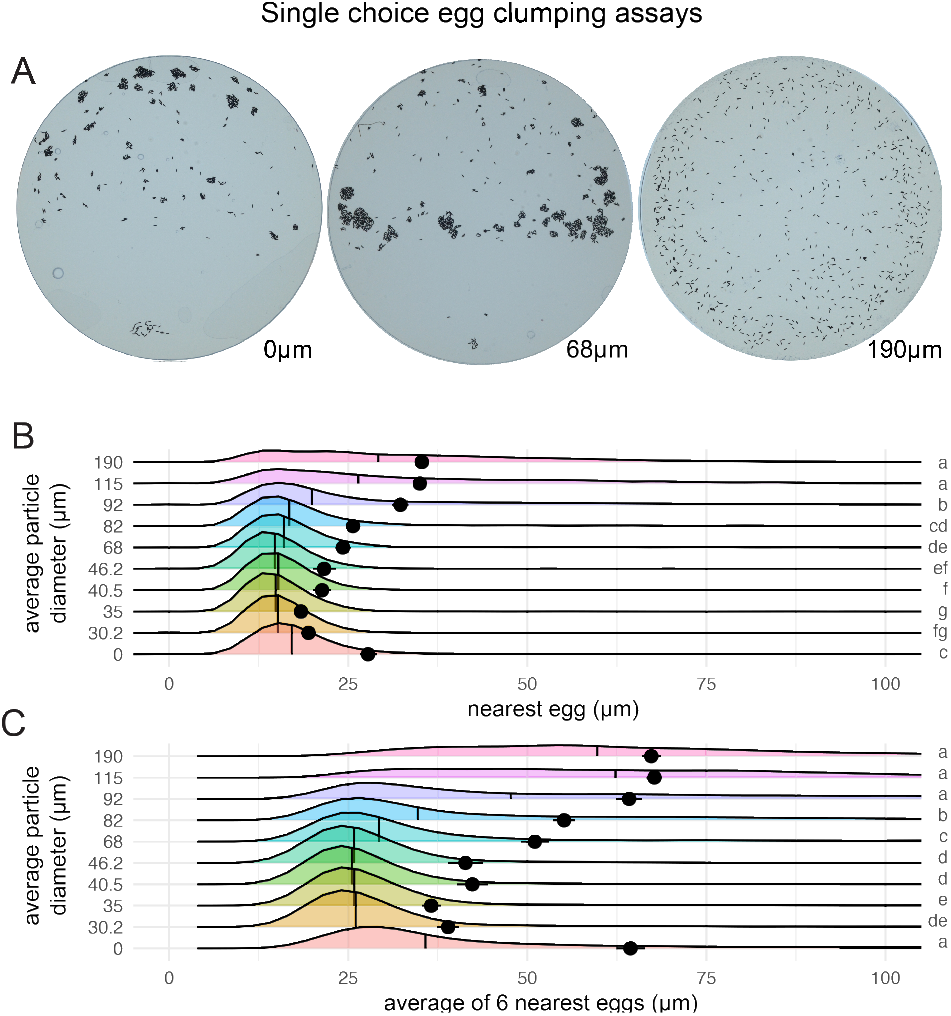
Eggs are laid in clumps on smooth surfaces. **(A)** Representative plates from plates with the indicated texture (average particle diameter) reveal differential levels of ‘clumping’ across textures. Quantification of the spatial arrangement of eggs relative to **(B)** their single nearest-neighbor **(C)** and average of the six nearest-neighbors. For B and C, vertical line indicates the median of the distribution, while the dot and whiskers represent mean and 95% confidence intervals. Letters indicate significant differences between group means as determined by one-way ANOVA followed by Tukey HSD pairwise tests. n=2109-4402 eggs analyzed per condition.

### Interaction of substrate texture and water salinity

We next wondered whether the presence of an aversive egg-laying cue (elevated salt) would influence texture preference in *Ae. aegypti* mosquitoes. We first repeated our split-dish assays but instead of providing fresh water, we provided a solution of 200mOsm/kg NaCl, which is an aversive cue (*11*). We found that the preference for rougher textures remained in the presence of salt (Figure 4A) and in fact was nearly identical to a similar curve generated in fresh water (Figure 4B), suggesting that mosquitoes can evaluate the relative value of multimodal cues independently when determining ideal egg-laying sites. To test the interaction between salt (a negative egg-laying cue) and rough texture (a positive egg-laying cue), we set up two-choice assays between smooth and salt-free substrates and rough and salted substrates at two different concentrations. We found a significant effect of salt concentration on egg-laying preference (Figure 4C), with the higher concentration of salt completely abolishing the positive preference for rough textures. The degree to which salt reduced the preference for rough textures did not differ with roughness (Figure 4D). This indicates that mosquitoes can integrate two competing cues, but that at higher concentrations of salt, texture is not relevant.

**Figure 4.**
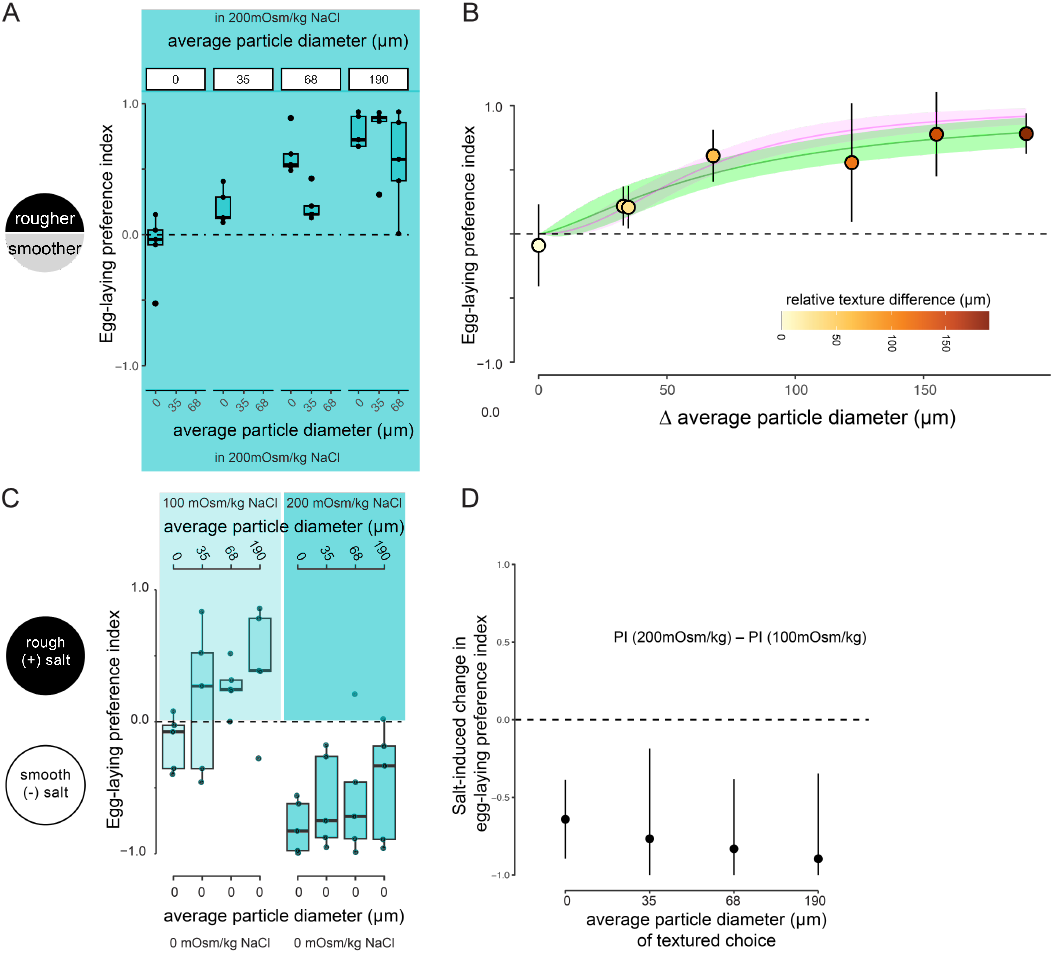
Texture and salt are interacting egg-laying cues. **(A)** Split-dish texture assays performed with 200mOsm/kg NaCl water for both choices. **(B)** Log-logistic function (fixed upper bound of 1 and lower bound of 0) fit to the egg-laying preference data shown in A (slope (b) = −1.407; ED50 (e) = 73.63) shown in green; log logistic function fit from experiments with no salt in Figure 2B overlaid in magenta **(C)** Two-choice egg-laying assays comparing preference for smooth surfaces with fresh water and rough surfaces with salt water at 100 mOsm/kg (left) or 200 mOsm/kg (right) respectively. A two-way ANOVA (PI ∼ salt * texture) revealed a significant effect of salt (p=2.9×10^−7^) **(D)** The difference (mean and 95% confidence intervals) in preference index in 200mOsm/kg NaCl – 100mOsm/kg NaCl. n=5 per condition (A) and n=5 per condition, (C).

### Surface texture changes locomotor behaviour egg-laying

We next asked whether surface texture played a role in movement dynamics during mosquito egg-laying by filming individual mosquitoes on a split-petri dish assay in which one half was smooth agarose and the other was textured with 190µm sandpaper (Figure 5A). We tracked position and manually scored egg-laying events for nine mosquitoes over 120 minutes and found that mosquitoes routinely traversed the boundary between smooth and textured sides of the plate, a sign of active exploration and local search during egg-laying (Figure 5B-C).

**Figure 5.**
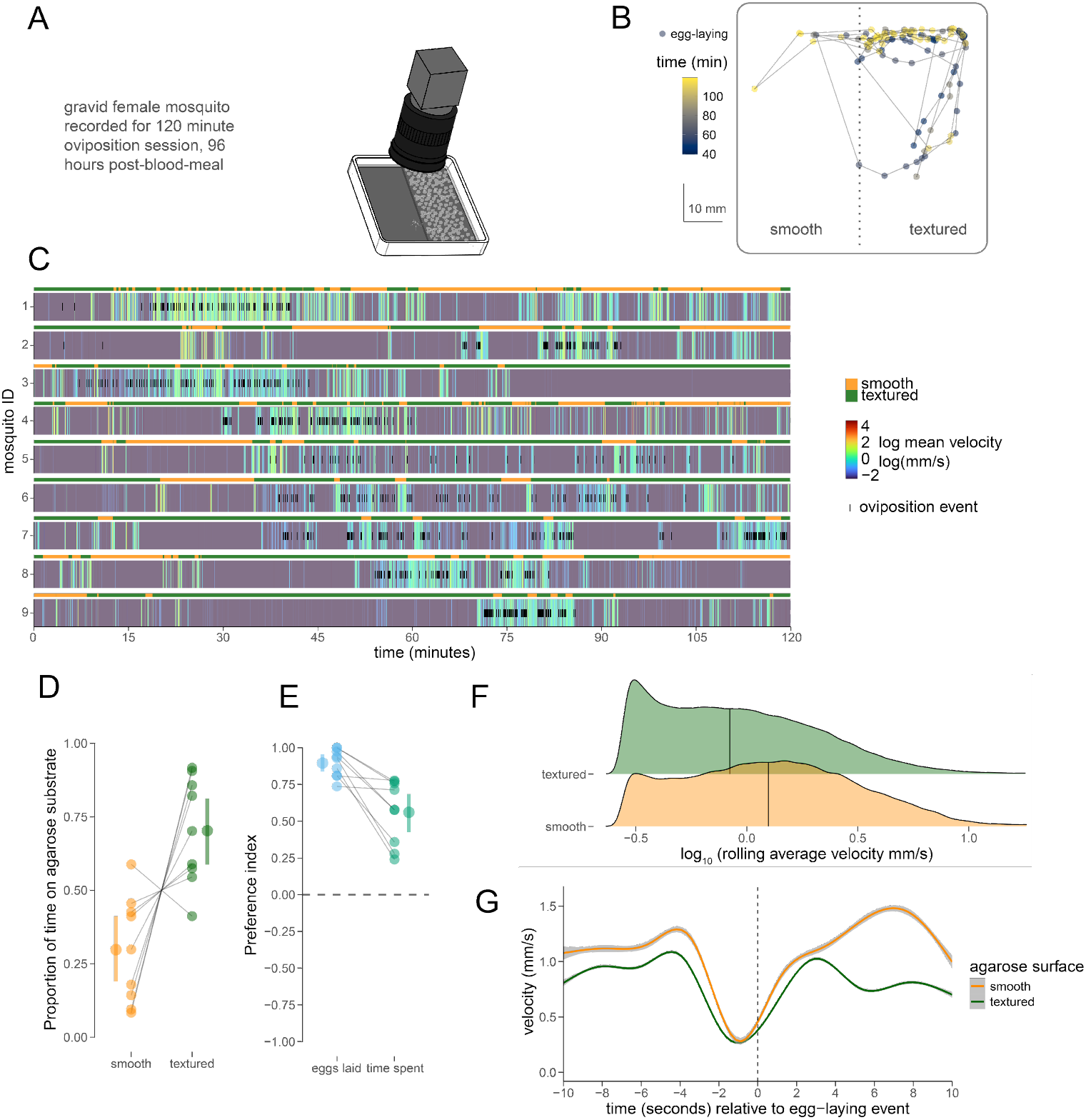
Substrate texture shapes mosquito movement during egg-laying. **(A)** Schematic of the behavioral arena and multi-camera tracking setup used to record individual gravid female mosquitoes during a 120-min egg-laying session (96 h post-blood meal). Females were allowed to move freely between smooth and textured agarose substrates. **(B)** Example trajectories of a representative individual overlaid on the arena, with positions colored by instantaneous velocity. **(C)** Raster plot summarizing behavioral state and position across time for individual mosquitoes (rows) during the recording session. Colored segments indicate movement speed and substrate occupancy over time. **(D)** Proportion of time spent on smooth versus textured substrates for individual mosquitoes, shown as paired measurements. **(E)** Relationship between time spent on each substrate and the number of eggs laid, shown for individual animals. **(F)** Distribution of rolling-average locomotor velocities on smooth and textured substrates, plotted on a log_10_ scale. Vertical line represents median **(G)** Average velocity profiles (rolling average of velocity with 0.5s window; mean and 95% confidence intervals) aligned to egg-laying events, shown separately for smooth and textured agarose. n=9 recordings.

We found that mosquitoes spent significantly more time on the textured half of the plate (Figure 5D) but, when expressed as a preference index, their place preference was significantly less than their egg-laying preference (Figure 5E). This indicates that egg-laying preference reflects an increased probability of laying eggs on rough textures and is not simply a product of preferring to spend time on rough surfaces. When examining a mosquito’s velocity on each half of the plate we found that they moved significantly faster on smooth agarose (Figure 5F). When examining movement dynamics locked to egg-laying events we found that mosquitoes showed a characteristic ‘pause’ ∼1 second prior to the appearance of an egg followed by a rebound in velocity that was significantly higher when animals were on smooth as compared to textured substrate (Figure 5G). Together, this suggests that mosquitoes behave differently on different textured surfaces.

## Discussion

### The importance of mechanical cues in egg-laying decisions

Our results highlight the role of interacting cues in dictating egg-laying site selection. While some cues guide mosquitoes to bodies of water from a distance, contact-mediated evaluation of water and substrate quality represents a final ‘go / no-go’ decision checkpoint prior to egg deposition. In this framework, evaluation of water quality (e.g. for appropriate salinity and absence of predators) unlocks a final search behaviour which culminates in the identification of a particular patch of surface on which to affix each egg.

By choosing textured surfaces, the female mosquito may be maximizing surface area of contact between egg and substrate, which could reduce the likelihood of an egg becoming prematurely dislodged before submergence. Future experiments to determine the force required to dislodge an egg on a smooth vs. a textured surface could directly demonstrate this relationship. In addition, depositing eggs in grooves and other features may buffer them from changes in temperature or humidity (*23*) or provide protection from predators (*24, 25*). Overall, ensuring adherence to substrate and protecting them from the elements could be critically important for the ability of *Ae. aegypti* mosquitoes to repopulate areas of their range which experience strong seasonality in precipitation, temperature, or other climatic factors.

In other species, mechanical properties of a particular substrate can influence egg-laying decisions. For example, the spotted wing Drosophila *Dropsophila suzukii* appears to lay eggs in a curvature-dependent fashion (*26*)

We found that mechanical (texture) and chemical cues (salinity) interact in a hierarchical fashion to guide an overall egg-laying decisions in *Ae. aegypti*. This is true in other insects as well, including the *D. suzukii* where the effect of surface hardness can supersede a preference against chemical cues derived from microbes (*27*). Future studies on egg-laying preference should thus consider the potential for combinatorial or hierarchical interactions between egg-laying cues, positive and negative.

### The cellular and molecular basis of texture discrimination during egg-laying

As the mosquito pauses just before laying an egg, we posit that a close evaluation of surface texture (likely in conjunction with other cues including surface moisture and hardness) guides the selection of a specific location for each egg. This final search likely involves deflection of mechanosensory hairs on the ovipositor of the mosquito (*28*) which would allow them to identify appropriately sized divots or other features in which to deposit each egg. In *Drosophila*, labellar and tarsal mechanoreceptors also appear to be involved in texture and stiffness discrimination during egg-laying (*29*). This raises the possibility that mosquitoes use sensory appendages beyond the ovipositor to evaluate surface texture.

Mechanosensation in insects relies on receptors from several gene families including piezo, ppk, Tmc, and TRP channels (*30*). The mosquito ovipositor and tarsi express members of each of these gene families (*31–33*), and in other species, have been implicated in texture-drive egg-laying behaviour (*29, 34–36*). Identification of mechanosensory receptors and cell types responsible for egg-laying site selection is an important goal of future research.

### Relevance to disease transmission

The yellow fever mosquito *Ae. aegypti* is an important vector of arboviral pathogens that cause dengue fever, chikungunya, Zika, yellow fever, and other diseases. Combined, these diseases kill tens of thousands of people every year. Pathogen acquisition by vector mosquitoes requires multiple blood-meals, and the link between egg-laying and a release of blood-meal induced host-seeking suppression in some mosquitoes, including *Ae. aegypti*, makes egg-laying an important component of arboviral disease spread (*37*). Furthermore, the search for egg-laying sites appears to drive dispersal of mosquitoes post-blood meal (*38*). In addition, identifying cues that guide egg-laying can help build better tools for mosquito surveillance and control, including gravid ovitraps (*18, 39*).

### Comparative considerations

There are over 3600 species of mosquito, many of which exhibit unique egg-laying strategies. These include species that eggs directly on water in rafts, those that deposit eggs deep into moist soil substrates where they can remain for years, or even those that deposit eggs aerially (e.g. *Sabethes*) (*40*). It will be important to examine the relative role of mechanosensation in egg-laying behaviours across species, with potential implications for understanding the limits of spread for different species.

## Acknowledgements

We would like to thank Jean-François Doherty, Sarah Garcia, Elva Vidya, and members of the Matthews Lab for comments on this manuscript and feedback throughout the course of the project. This work was supported by grants from Natural Sciences and Engineering Research Council of Canada (NSERC Discovery Grant RGPIN-2020-05423 to BJM; NSERC Undergraduate Student Research Award to AA), Canadian Institutes of Health Research (CIHR Project Grant PJT-191985 to BJM), Michael Smith Health Research BC (MSHRBC Scholar Award SCH-2021-1860 to BJM), Alfred P. Sloan Foundation (Sloan Research Fellowship in Neuroscience FG-2021-16383 to BJM), and Human Frontier Science Program (HFSP Research Grant RGP018/2023 to BJM). Additional infrastructure support was provided by the Canadian Foundation for Innovation and British Columbia Knowledge Development Fund (CFI JELF and BCKDF awards 40036, 41223, 43348 to BJM) and unrestricted startup funds from The University of British Columbia (BJM). This paper was typeset with a template: www.github.com/chrelli/bioRxiv-word-template

## Author contributions

Study design and conceptualization: AA, BJM. Development of methodology: AA, NT, AZ, LS, BJM Data acquisition: AA, NT, AZ, LS

Data analysis, curation, and visualization: AA, NT, AZ, LS, BJM Writing, original draft preparation: AA, BJM

Writing, review and editing: AA, NT, AZ, LS, BJM

Supervision, funding acquisition, and project administration: BJM

## Materials and Methods

### Experimental Model and Organisms

*Aedes aegypti* mosquitoes (Orlando strain; Matthews lab stock) were reared and maintained in an insectary at 25-28°C, 70-80% relative humidity with a 14-10 light-dark cycle. Mosquito eggs were hatched under vacuum for 30 minutes and reared in dechlorinated Vancouver tap water at densities of 250 larvae per 2-2.5 L of water and temperature of 27 C while being fed ground fish flakes (TetraMim Tropical Flakes, Tetra) as needed.

Pupae were transferred to 30 cm^3 Bugdorm cages at densities of 300-350 animals per cage. Adults were provided 10% sucrose ad libitum for 9-13 days before being blood-fed with defibrinated sheep’s blood (Hardy Diagnostics) in parafilm-covered glass feeders (Chemglass, 50 mm diameter) with 37 °C water circulating through an outer water jacket or a Hemotek blood-feeding apparatus to maintain temperature. After blood-feeding, engorged females were visually selected and transferred to a new cage with sucrose and used for behaviour experiments 4 days after the blood meal. Females were not used beyond their first gonotrophic (egg-laying) cycle.

### Preparation of textured agarose plates

A 3% w/v molten agarose solution (BioBasic, Agarose A) was made by dissolving 3 grams powder per 100 mL of dechlorinated water and microwaved until all agarose was dissolved and the solution was bubbling. Agarose was cooled at room temperature until warm to touch (∼60°C). Large petri dishes (15 mm diameter, Fisher Scientific) were lined with sandpaper of desired grit (3M, ranging 30.2 µm to 190 µm average particle size; see table S1). Sandpaper disks were cut and glued to the dishes textured side up. 75 mL of warm 3% agarose solution was poured and allowed to set until firm. Circles of clear, solidified agarose were cut with a sterile blade and transferred to smaller 60mm petri dishes (Fisher Scientific) with the textured side up. The same procedure was repeated for smooth plates but using the reverse (smooth) side of a sandpaper disk to imprint a flat surface.

### Two-choice egg-laying assays

White restaurant supply buckets (85oz volume) with a fine mesh top covering were used as arenas for egg-laying assays. Two petri dishes (60mm in diameter) were mounted on angled support structures (tilted at ∼20°) facing towards the center of the arena, 1-2 cm apart. 2 mL of dechlorinated tap water was pooled at the bottom of each dish. 10 gravid females were added to each arena, which was then placed in the insectary for 22-24 hours at 70-80% humidity and 25-28° C. Placement of texture treatments was randomized within each arena and the arenas were distributed randomly in the insectary.

### Photographing of egg papers

Eggs were retrieved on petri dishes after 22-24 hours of egg-laying and counted under a dissecting scope by hand using a spatula to move the eggs as they were counted. Raw egg counts were recorded for each pair of dishes. For automated counting algorithms, dishes were photographed using an SLR camera (Nikon D7000) with a macro lens (Tamron 90mm F/2.8 Di SP MACRO 1:1) at a set height of 44 cm. Dishes were illuminated using an LED backlight to enhance contrast of melanized eggs against agarose. Images were cropped using Adobe Photoshop CS6 to form 3200 x 3200 pixel-sized circles of the inner portion of the dish along the edges of the agarose substrate.

## Data analysis

Eggs were manually counted and a preference index was calculated for each two-choice assay using the following formula:

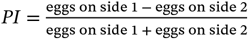

For split-dish two-choice assays (Figure 2), eggs within 3mm of the midline (where the two textures met) were not considered in calculating the preference index. The data was visualized with R (*41*) and RStudio (*42*) using the following packages: ggplot2 (*43*), drc (*44*), and tidyverse (*45*). Details of specific visual representations and statistical tests used are reported in the appropriate figure legends.

For clumping analysis, a separate two-choice assay experiment was repeated using the setup described above. Representative, high-resolution images of dishes of tested textures were photographed an SLR camera (Nikon D7000) with a macro lens (Tamron 90mm F/2.8 Di SP MACRO 1:1) at a set height of 44 cm and analyzed for egg clustering using FIJI/ImageJ (*46*). A circular image was turned to an 8-bit image. Egg centroids were first selected with Process > Find Maxima… with “Prominence” set above 100.00, the “Light background” box checked, and “Point selection” as the output type. Then, the multi-point selection tool was used to go over the image manually to select any non-selected centroids and delete erroneous selections of non-egg particles. Once all centroids were selected, a macro https://imagejdocu.tudor.lu/macro/multiple_points was used to export the points into .csv and excel file formats. A complementary macro from the same source was used to import the points back onto the image.

To determine the average distance to the six closest neighbours of each centroid, a plugin was used as described (*47*). With points selected on the image, Analyze > Measure was used followed by the ND plugin with “6” as the number of neighbours.

### Imaging and analysis of temporal dynamics

For live imaging experiments, mosquitoes were reared as above and bloodfed 96 hours prior to each assay. Fully engorged mosquitoes were selected and provided sucrose ad libitum prior to recording. Imaging sessions took place during the mosquito’s subjective day (ZT +6, +9, or +12 hours of a 14:10 light:dark cycle).

Mosquitoes were provided 3% agarose as prepared above, first by preparing a 150mm petri dish with equally sized smooth and rough (190µm average particle size) halves. From this, a 65mm^2^ rectangular slab was carved with a custom 3D printed cutter, with equally sized smooth and rough halves. Agarose slabs were stored in dechlorinated water until use.

For each imaging session, a single mosquito was cold-anesthetized and then placed into a small arena with the agarose slab at its center, tilted ∼20°. 2mL of dechlorinated tap water was added to the bottom of this slanted arena to stimulate egg-laying. The lid to a 90mm petri dish was placed over the arena to contain the mosquito, and the entire apparatus was contained within a blackout box (no visible light) and the arena was illuminated with infrared LEDs (940nm). A 120-minute recording was triggered using the GPIO pins on a Basler a2A2448-75umPRO camera with a Marshall Electronics 11-50mm f1.4 cs-mount camera lens (CS1150-8MP) with a 3D-printed aperture insert with a constant 3 mm diameter.

Centroid tracking of mosquitoes in each video was accomplished using SLEAP v1.4.2 (*48*) and egg-laying events were manually scored. A model skeleton of the mosquito was produced in an iterative process alternating between manual training and automated model refinement. All point data were calculated using the centroid of the mosquito’s thorax, which was considered the centroid of the mosquito for tracking purposes. These centroid coordinates were exported as a csv and processed using R v4.4.1 (*41*) for further analysis. Position of the mosquito and each egg was calculated relative to the midline, and a rolling average of velocity was calculated with a window of 0.5s.

## References

1. M. D. Bentley, J. F. Day, Chemical Ecology and Behavioral Aspects of Mosquito Oviposition. Annu. Rev. Entomol. 34, 401–421 (1989).

2. J. F. Day, Mosquito Oviposition Behavior and Vector Control. Insects 7, 65 (2016).

3. S. R. Christophers, Aedes Aegypti (L.) the Yellow Fever Mosquito: Its Life History, Bionomics and Structure (Cambridge University Press, Cambridge, 2009).

4. M. Mayilsamy, Extremely Long Viability of Aedes aegypti (Diptera: Culicidae) Eggs Stored Under Normal Room Condition. J. Med. Entomol. 56, 878–880 (2019).

5. J. R. Powell, A. Gloria-Soria, P. Kotsakiozi, Recent History of Aedes aegypti: Vector Genomics and Epidemiology Records. Bioscience 68, 854–860 (2018).

6. N. H. Rose, M. Sylla, A. Badolo, J. Lutomiah, D. Ayala, O. B. Aribodor, N. Ibe, J. Akorli, S. Otoo, J.-P. Mutebi, A. L. Kriete, E. G. Ewing, R. Sang, A. Gloria-Soria, J. R. Powell, R. E. Baker, B. J. White, J. E. Crawford, C. S. McBride, Climate and urbanization drive mosquito preference for humans. Curr. Biol. 30, 3570–3579 (2020).

7. M. U. G. Kraemer, R. C. Reiner, O. J. Brady, J. P. Messina, M. Gilbert, M. Pigott, D. Yi, K. Johnson, L. Earl, L. B. Marczak, S. Shirude, N. Davis Weaver, D. Bisanzio, T. A. Perkins, S. Lai, X. Lu, P. Jones, G. Coelho, R. G. Carvalho, W. Van Bortel, C. Marsboom, G. Hendrickx, F. Schaffner, C. G. Moore, H. H. Nax, L. Bengtsson, E. Wetter, A. J. Tatem, J. S. Brownstein, D. L. Smith, L. Lambrechts, S. Cauchemez, C. Linard, N. R. Faria, O. G. Pybus, T. W. Scott, Q. Liu, H. Yu, G. R. W. Wint, S. I. Hay, N. Golding, Past and future spread of the arbovirus vectors Aedes aegypti and Aedes albopictus. Nat. Microbiol. 4, 854–863 (2019).

8. M. U. G. Kraemer, M. E. Sinka, K. A. Duda, A. Q. N. Mylne, F. M. Shearer, C. M. Barker, C. G. Moore, R. G. Carvalho, G. E. Coelho, W. Van Bortel, G. Hendrickx, F. Schaffner, I. R. F. Elyazar, H.-J. Teng, O. J. Brady, J. P. Messina, D. M. Pigott, T. W. Scott, D. L. Smith, G. R. W. Wint, N. Golding, S. I. Hay, The global distribution of the arbovirus vectors Aedes aegypti and Ae. albopictus. eLife 4, e08347 (2015).

9. W. J. Laursen, G. Budelli, R. Tang, E. C. Chang, R. Busby, S. Shankar, R. Gerber, C. Greppi, R. Albuquerque, P. A. Garrity, Humidity sensors that alert mosquitoes to nearby hosts and egg-laying sites. Neuron, S0896627322011229 (2023).

10. M. Bar-Zeev, Oviposition of Aedes aegypti L. on a dry surface and hygroreceptors. Nature 213, 737–738 (1967).

11. B. J. Matthews, M. A. Younger, L. B. Vosshall, The ion channel ppk301 controls freshwater egg-laying in the mosquito Aedes aegypti. eLife 8, e43963 (2019).

12. A. Afify, C. G. Galizia, Chemosensory Cues for Mosquito Oviposition Site Selection. J. Med. Entomol. 52, 120–130 (2015).

13. A. O’Gower, The influence of the surface on opposition by Aedes aegypti (Linn.)(Diptera, Culicidae). Proc Linn. Soc New South Wales 82, 240–244 (1957).

14. J. C. Rojas, A. Virgen, L. Cruz-López, Chemical and Tactile Cues Influencing Oviposition of a Generalist Moth, <I>Spodoptera frugiperda</I> (Lepidoptera: Noctuidae). Environ. Entomol. 32, 1386– 1392 (2003).

15. J. A. A. Renwick, F. S. Chew, Oviposition Behavior in Lepidoptera. Annu. Rev. Entomol. 39, 377–400 (1994).

16. W. Beckel, Oviposition site preference of Aedes mosquitoes (Culicidae) in the laboratory. Mosq. News 15, 224–228 (1955).

17. R. C. Wallis, Observations on Oviposition of Two Aedes Mosquitoes (Diptera Culicidae)1. Ann. Entomol. Soc. Am. 47, 393–396 (1954).

18. M. Momen, K. Seheli, Md. A. Hossain, A. Ghosh, Md. F. Hossain, Effect of different lining paper materials and infusions on oviposition preference of Aedes aegypti (Diptera: Culicidae) gravid mosquitoes. Sci. Rep. 15, 16379 (2025).

19. D. D. Chadee, P. S. Corbet, H. Talbot, Proportions of eggs laid by Aedes aegypti on different substrates within an ovitrap in Trinidad, West Indies. Med. Vet. Entomol. 9, 66–70 (1995).

20. R. Fay, A. Perry, Laboratory studies of ovipositional preferences of Aedes aegypti. Mosq. News 25, 276–281 (1965).

21. J. Wong, H. Astete, A. C. Morrison, T. W. Scott, Sampling considerations for designing Aedes aegypti (Diptera: Culicidae) oviposition studies in Iquitos, Peru: substrate preference, diurnal periodicity, and gonotrophic cycle length. J. Med. Entomol. 48, 45 (2011).

22. A. K. O’Gower, Environmental stimuli and the oviposition behaviour of Aedes aegypti var. queenslandis Theobald (Diptera, Culicidae). Anim. Behav. 11, 189–197 (1963).

23. R. Russo, Substrate Texture as an Oviposition Stimulus for Aedes Vexans (Diptera: Culicidae). J. Med. Entomol. 15, 17–20 (1978).

24. H. James, Location of univoltine Aedes eggs in woodland pool areas and experimental exposure to predators. Mosq. News 26, 59–63 (1966).

25. B. Byttebier, S. Fischer, Predation on Eggs of Aedes aegypti (Diptera: Culicidae): Temporal Dynamics and Identification of Potential Predators During the Winter Season in a Temperate Region. J. Med. Entomol. 56, 737–743 (2019).

26. J. Akutsu, T. Matsuo, Drosophila suzukii preferentially lays eggs on spherical surfaces with a smaller radius. Sci. Rep. 12, 15792 (2022).

27. A. Sato, K. M. Tanaka, J. Y. Yew, A. Takahashi, Drosophila suzukii avoidance of microbes in oviposition choice. R. Soc. Open Sci. 8, 201601 (2021).

28. P. A. Rossignol, S. B. McIver, Fine structure and role in behavior of sensilla on the terminalia of Aedes aegypti (L.) (Diptera: Culicidae). J. Morphol. 151, 419–437 (1977).

29. L. Zhang, J. Yu, X. Guo, J. Wei, T. Liu, W. Zhang, Parallel Mechanosensory Pathways Direct Oviposition Decision-Making in Drosophila. Curr. Biol. 30, 3075-3088.e4 (2020).

30. P. Hehlert, W. Zhang, M. C. Göpfert, Drosophila Mechanosensory Transduction. Trends Neurosci. 44, 323–335 (2021).

31. B. J. Matthews, O. Dudchenko, S. B. Kingan, S. Koren, I. Antoshechkin, J. E. Crawford, W. J. Glassford, M. Herre, S. N. Redmond, N. H. Rose, G. D. Weedall, Y. Wu, S. S. Batra, C. A. Brito-Sierra, S. D. Buckingham, C. L. Campbell, S. Chan, E. Cox, B. R. Evans, T. Fansiri, I. Filipović, A. Fontaine, A. Gloria-Soria, R. Hall, V.S. Joardar, A. K. Jones, R. G. G. Kay, V. K. Kodali, J. Lee, G. J. Lycett, S. N. Mitchell, J. Muehling, M. R. Murphy, A. D. Omer, F. A. Partridge, P. Peluso, A. P. Aiden, V. Ramasamy, G. Rašić, S. Roy, K. Saavedra-Rodriguez, S. Sharan, A. Sharma, M. L. Smith, J. Turner, A.M. Weakley, Z. Zhao, O. S. Akbari, W. C. Black, H. Cao, A. C. Darby, C. A. Hill, J. S. Johnston, T. D. Murphy, A. S. Raikhel, D. B. Sattelle, I. V. Sharakhov, B. J. White, L. Zhao, E. L. Aiden, R. S. Mann, L. Lambrechts, J. R. Powell, M. V. Sharakhova, Z. Tu, H. M. Robertson, C. S. McBride, A. R. Hastie, J. Korlach, D. E. Neafsey, A. M. Phillippy, L. B. Vosshall, Improved reference genome of Aedes aegypti informs arbovirus vector control. Nature 563, 501–507 (2018).

32. B. J. Matthews, C. S. McBride, M. DeGennaro, O. Despo, L. B. Vosshall, The neurotranscriptome of the Aedes aegypti mosquito. BMC Genomics 17, 32 (2016).

33. O. V. Goldman, A. E. DeFoe, Y. Qi, Y. Jiao, S.-C. Weng, B. Wick, L. Houri-Zeevi, P. Lakhiani, T. Morita, J. Razzauti, A. Rosas-Villegas, Y. N. Tsitohay, M. M. Walker, B. R. Hopkins, J. X. D. Ang, I. Antoshechkin, Y. Cai, F. Chen, Y.-C. Chen, J. Devilliers, L. Dong, R. Feuda, P. Gabrieli, A. Kopp, H. Kwon, H.-H. Li, T.-C. Lu, T. Lucio, J. T. Marques, M. F. Oliveira, R. P. Olmo, U. Palatini, Z. M. Pithawala, J. Pompon, Y. Reis, J. Rodrigues, R. C. Smith, M. Haeussler, O.S. Akbari, L. B. Duvall, H. White-Cooper, T. R. Sorrells, R. Sharma, H. Li, L. B. Vosshall, N. Shai, A single-nucleus transcriptomic atlas of the adult Aedes aegypti mosquito. Cell, S0092867425011377 (2025).

34. S.-F. Wu, Y.-L. Ja, Y. Zhang, C.-H. Yang, Sweet neurons inhibit texture discrimination by signaling TMC-expressing mechanosensitive neurons in Drosophila. eLife 8, e46165 (2019).

35. L. Guo, Z.-D. Zhou, F. Mao, X.-Y. Fan, G.-Y. Liu, J. Huang, X.-M. Qiao, Identification of potential mechanosensitive ion channels involved in texture discrimination during Drosophila suzukii egg-laying behaviour. Insect Mol. Biol. 29, 444–451 (2020).

36. G. Zhang, C. Wu, X. Cui, H. Wang, B. Wei, A. Mazarin, A. Mansour, C. Niu, Multiple mechanosensory pathways mediate oviposition behavior in Bactrocera dorsalis (Diptera: Tephritidae). Pest Manag. Sci. 81, 5551–5560 (2025).

37. L. B. Duvall, Mosquito Host-Seeking Regulation: Targets for Behavioral Control. Trends Parasitol. 35, 704–714 (2019).

38. P. Reiter, M. A. Amador, R. A. Anderson, G. G. Clark, Short Report: Dispersal of Aedes aegypti in an Urban Area after Blood Feeding as Demonstrated by Rubidium-Marked Eggs. Am. J. Trop. Med. Hyg. 52, 177–179 (1995).

39. P. A.P.S., Optimizing ovitrap design: the role of ovistrip texture, colour, and water in modulating oviposition behavior of Aedes vector mosquitoes. Trop. Biomed. 42, 459–466 (2025).

40. G. Vieira, M. I. L. Bersot, G. R. Pereira, F. V. S. de Abreu, A. C. Nascimento-Pereira, M. S. A. S. Neves, M. G. Rosa-Freitas, M. de A. Motta, R. Lourenco-de-Oliveira, “Oviposition in flight: the Sabethes albiprivus incredible egg throwing behavior” (preprint, Zoology, 2020); 10.1101/2020.01.31.927228.

41. R Core Team, R: A Language and Environment for Statistical Computing (R Foundation for Statistical Computing, Vienna, Austria, 2025; https://www.R-project.org/).

42. Posit team, RStudio: Integrated Development Environment for R, Posit Software, PBC (2025).

43. H. Wickham, Ggplot2: Elegant Graphics for Data Analysis (Springer International Publishing, Cham, Switzerland, ed. 2, 2016) Use R!

44. C. Ritz, F. Baty, J. C. Streibig, D. Gerhard, Dose-Response Analysis Using R. PLOS ONE 10, e0146021 (2015).

45. H. Wickham, M. Averick, J. Bryan, W. Chang, L. McGowan, R. François, G. Grolemund, A. Hayes, L. Henry, J. Hester, M. Kuhn, T. Pedersen, E. Miller, S. Bache, K. Müller, J. Ooms, D. Robinson, D. Seidel, V. Spinu, K. Takahashi, D. Vaughan, C. Wilke, K. Woo, H. Yutani, Welcome to the tidyverse. J Open Source Softw 4, 1686 (2019).

46. J. Schindelin, I. Arganda-Carreras, E. Frise, V. Kaynig, M. Longair, T. Pietzsch, S. Preibisch, C. Rueden, S. Saalfeld, B. Schmid, J.-Y. Tinevez, D. J. White, V. Hartenstein, K. Eliceiri, P. Tomancak, A. Cardona, Fiji: an open-source platform for biological-image analysis. Nat. Methods 9, 676–682 (2012).

47. M. Haeri, M. Haeri, ImageJ Plugin for Analysis of Porous Scaffolds used in Tissue Engineering. J. Open Res. Softw. 3 (2015).

48. T. D. Pereira, N. Tabris, A. Matsliah, D. M. Turner, J. Li, S. Ravindranath, E. S. Papadoyannis, E. Normand, D. S. Deutsch, Z. Y. Wang, G. C. McKenzie-Smith, C. C. Mitelut, M. D. Castro, J. D’Uva, M. Kislin, D. H. Sanes, S. D. Kocher, S. S.-H. Wang, A. L. Falkner, J. W. Shaevitz, M. Murthy, SLEAP: A deep learning system for multianimal pose tracking. Nat. Methods 19, 486–495 (2022).

